# larch: mapping the parsimony-optimal landscape of trees for directed exploration

**DOI:** 10.1101/2025.10.29.685337

**Authors:** Mary Barker, Ognian Milanov, Will Dumm, Dave Rich, Yatish Turakhia, Frederick A Matsen

**Affiliations:** Computational Biology Program, Fred Hutchinson Cancer Research Center, Seattle, Washington, USA; Howard Hughes Medical Institute, Computational Biology Program, Fred Hutchinson Cancer Research Center, Seattle, Washington, USA; Department of Electrical and Computer Engineering, University of California San Diego, San Diego, California, USA; Department of Genome Sciences, University of Washington, Seattle, Washington, USA; Department of Statistics, University of Washington, Seattle, Washington, USA

## Abstract

Phylogenetic inference algorithms for large data sets typically return a single tree. However, there are often many optimal trees, especially when sequence data is closely related. We develop a compact representation of large collections of maximally parsimonious *histories*—trees with mutations mapped onto tree edges. Our C++ implementation, `larch`, leverages this representation for a highly parallel search algorithm. The storage component uses our history DAG structure to compactly represent large families of optimal trees. The search algorithm integrates this storage with `matOptimize` for rapid tree optimization; the DAG structure allows us to accept thousands of conflicting tree rearrangements in parallel. The integration enables a new type of tree search: one that systematically maps out the collection of good trees, enabling moves that are directed away from the current set of optimal trees to cross valleys and increase the diversity of the set of optimal trees. It is able to identify more parsimonious trees than are found by other methods. We find diverse optimality landscapes for viral datasets, including many distinct plateaux. We also find that our implementation produces similar results whether using a variety of single starting trees or an ensemble of starting trees, indicating effective global optimization.

## Introduction

It has long been recognized that many phylogenetic trees can suitably explain a given sequence data set. For maximum parsimony, early authors noticed many equally parsimonious trees [1]. For maximum likelihood, tests have been developed to detect “significant” differences in likelihood [2, 3, 4], implicitly acknowledging that there are often multiple statistically indistinguishable trees. Bayesian analyses explicitly integrate over tree uncertainty, and posterior support is rarely small.

What is the landscape of this good set of trees? For maximum parsimony, Maddison [5] first described “islands” of maximum parsimony trees, and explored their structure. For Bayesian analyses, interesting structure has been found for macroevolutionary [6] and viral [7, 8] datasets, in which there are distinct clusters of trees separated by valleys.

However, finding that ensemble of good trees is hard. Finding a single parsimony or maximum-likelihood tree is already NP-hard [9, 10, 11], and finding the ensemble of optimal trees is necessarily harder. Typical Bayesian inference uses MCMC, sometimes requiring very long runs to fully explore posterior distributions [7, 8]. Systematic inference of Bayesian posterior distributions is possible but does not scale [12]. Thus, the field lacks a means of finding the collection of optimal trees for a large collection of sequences.

Some important ingredients for finding this set of optimal trees are now available. Namely, `matOptimize` [13] is software for tree search that optimizes an existing tree using an iterative, highly parallelized SPR search routine. This routine uses a partial Fitch algorithm to compute the parsimony criterion to evaluate proposed SPR moves and applies a random subset of the optimal moves to improve or at least maintain the parsimony score. The `matOptimize` software is built to accommodate pandemic-scale data by introducing an efficient storage structure called a mutation annotated tree (MAT) [14] (which has precedent in earlier work [15]). The MAT is an effective structure for densely sampled data, as it does not store entire sequence reconstructions at any internal or leaf nodes. Instead, a single “reference sequence” is stored for the root node, and each edge of the tree is annotated with the location and nucleotide changes where the parent and child nodes of that edge differ.

Another ingredient is the history directed acyclic graph (hDAG): a generalization of a phylogenetic tree with ancestral states that can represent large families of maximally parsimonious trees [16]. We will use the word “history” to refer to a phylogenetic tree with ancestral states mapped onto the internal nodes. The hDAG stores fully resolved sequences and topological split data at internal nodes to preserve parsimony results across the histories it represents.

Having these ingredients in hand raises a tantalizing opportunity: *could we systematically map out the space of optimal trees and develop a search algorithm that directs itself to new areas that need to be explored?*

In this paper, we introduce such a method of exploring the space of histories. The hDAG can compactly store very many such histories, but for large data we further require a MAT-like optimization. We combine the hDAG and MAT ideas into a structure called a mutation annotated directed acyclic graph (MADAG, Fig. 1).

**Figure 1:**
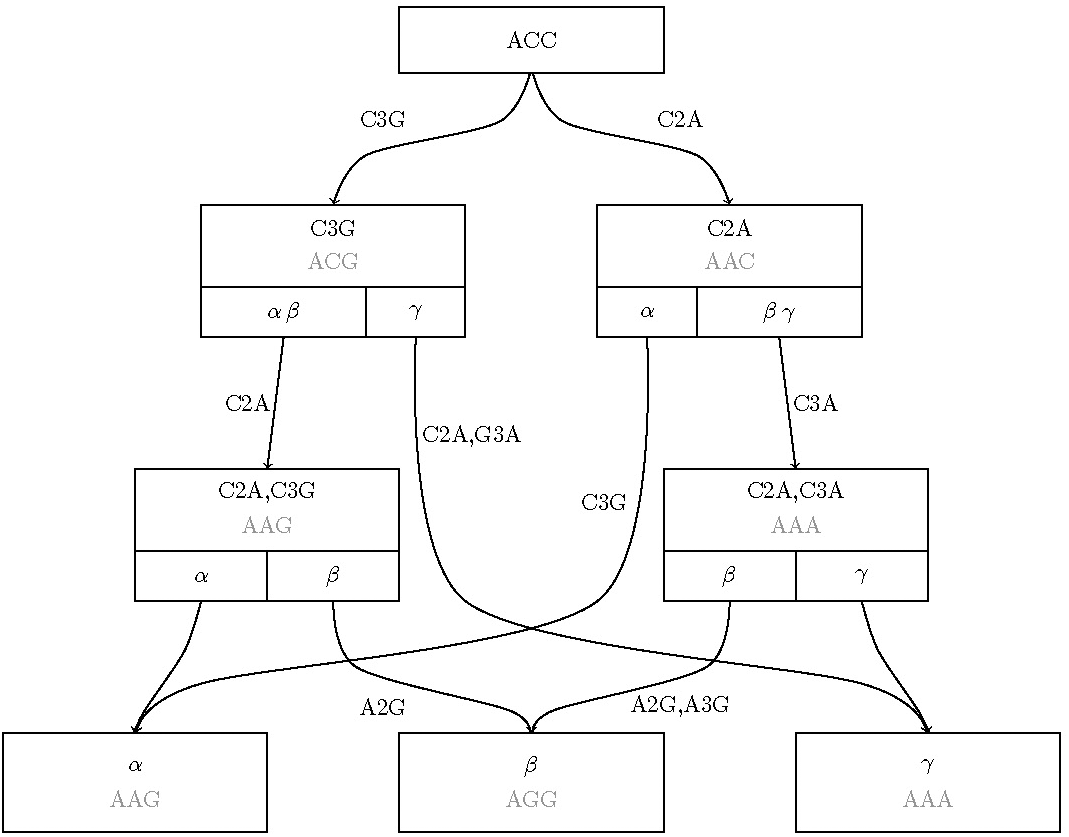
Example of a MADAG, which stores a full sequence at the root and uses edge mutation annotations for each edge and compact genome annotations at internal nodes. Nodes are also annotated with clade data, which consists of the partition of leaf nodes defined by their child edges. Sequence data for non-root nodes is shown in gray for readability, but is not stored directly in the MADAG.

This structure is equivalent in content to the hDAG; however, it uses the mutation-level edge annotation like the MAT. As with the hDAG, the MADAG reproduces the histories that are used to define it, but in general it can be used to recover many additional histories that are created by combining substructures from the original set.

We implement MADAG inference in `larch`, a new C++ program which achieves a highly efficient exploration of the space of optimal histories. The core of `larch` is an accelerated search routine based on `matOptimize` that integrates SPR moves directly into the MADAG. We are able to evaluate and apply thousands of SPR moves simultaneously and in parallel, even if they conflict. This feature is enabled by the MADAG and would not be possible with a treebased approach. The MADAG structure enables dynamic programs to store only maximally parsimonious histories, to sample histories from the MADAG based on a range of probability distributions, and to compute the Robinson-Foulds (RF) distances between each history and the remaining histories.

This allows us to develop new approaches for directed tree search, both in the way we sample histories with novel features and in the criterion used to filter the set of histories under consideration.

First, we find that incorporating topological novelty into the filtering objective function improves tree exploration. This topological novelty is calculated with respect to the entire currently known hDAG. By allowing a decrease in parsimony along the path to a new region, we find more histories with a given parsimony score, as well as histories that are themselves more optimal. We believe this is the first time that such a “valley-crossing” algorithm has been found that directly leverages knowledge of the optimality landscape.

Second, we find that applying SPR moves to topologically rare structures in the MADAG accelerates optimization. More specifically, the tree search process in `larch` iteratively samples histories on which SPRs are evaluated. We can guide the tree search to favor diversity by sampling the histories from the MADAG that are farthest from the medoid history. This outperforms other sampling methods, including sampling uniformly from the space of histories and sampling from strictly optimal histories.

These innovations work: we find that `larch` discovers more parsimonious individual trees than `matOptimize` and is able to quickly infer hDAGs with many histories in them. In a single iteration, we can build a MADAG with 588,825 leaves and 101,956,305,852 histories in 4 minutes, 10 seconds using a randomly-sampled subtree approach on the 20B dataset listed in Table 2. The same approach produces over 8.59 *×* 10^29^ histories in 5 iterations with a runtime of approximately 11 minutes.

Our method reveals the structure of the space of optimal trees. Specifically, we explore the optimality space for a variety of datasets, and show via Robinson-Foulds that we are finding topological diversity, not just node labeling diversity. We find clusters of optima that are topologically similar within each cluster yet distinct between clusters. We also demonstrate that `larch` achieves effective global exploration, as demonstrated by similar results from different starting points. Specifically, beginning with one tree gives essentially the same results as beginning with a collection of UShER-optimized trees with random taxon addition orders.

`larch` is available at https://github.com/matsengrp/larch/.

## Results

### Methods overview

The `larch` algorithm systematically explores maximally parsimonious history space while maintaining a compact ensemble representation. Here we give a brief overview of how this method works; full details can be found in the Methods section.

The `larch` algorithm can be described in general terms as a sample-optimizemerge loop. At each iteration, a history *T* is sampled from the MADAG and introduced to `matOptimize`. The SPR search of `matOptimize` uses a callback to rank the SPR moves that are found. Moves that satisfy parsimony or topological novelty criteria are approved and added to a list in order of their scores. The approved moves are also applied directly to the MADAG using a partial reconstruction of the altered subtree. At the end of each optimize step of `matOptimize`, a nonconflicting subset of the approved moves is chosen using a greedy algorithm and is applied to the sampled history, producing an optimized history that is then merged into the MADAG. This iterative process is described in Fig. 2.

**Figure 2:**
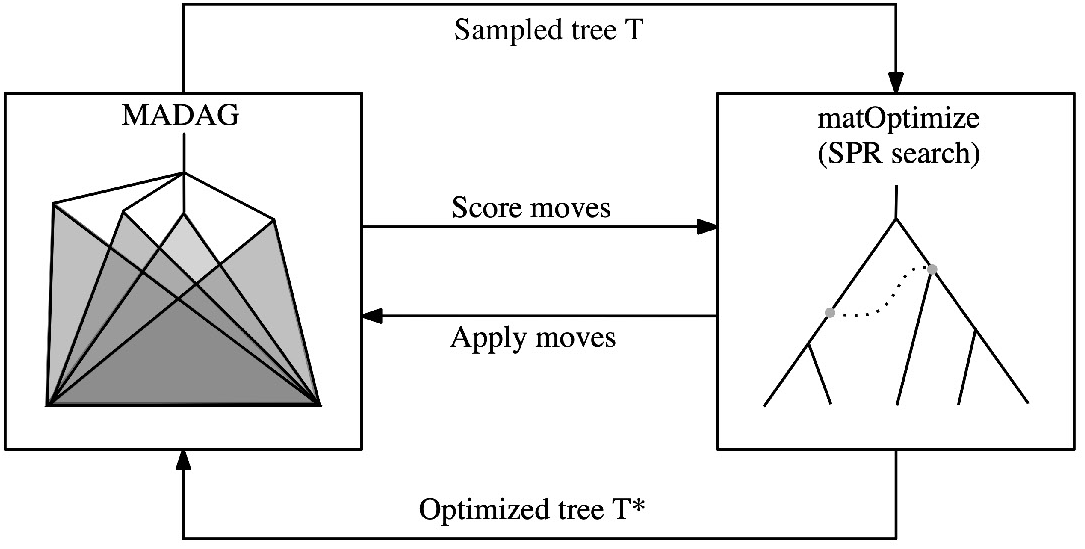
Workflow describing the sample-optimize-merge loop. Each `larch` iteration consists of sampling a single history from the MADAG and using it to identify and rank potential SPR moves using `matOptimize`. The optimal moves are applied directly to the MADAG, and a subset of the optimal moves are applied to the sampled history.

### Rewarding topological novelty improves tree search

Because we maintain a data structure that contains all the good trees, we can prioritize finding novel phylogenetic structures. We define <u>topological novelty</u> to be the number of new splits introduced to the MADAG by a proposed SPR move. We sought to understand if using topological novelty could improve hDAG inference.

In order to evaluate the effect of using different scoring criteria, we performed the following experiment: beginning with a single maximally parsimonious history, we considered two separate regimes for running `larch`. For the first instance, we selected the parsimony criterion to score moves (`score parsimony change`). With this criterion, SPR moves were ranked by the change in parsimony score. Only SPR moves with a score of 0 or lower, that is, scores that preserved or improved the parsimony score of the sampled history, were approved. (We note that the default behavior of `matOptimize` is to reject all moves that do not improve parsimony.) For the second instance, we used a combination of topological novelty and parsimony as a scoring criterion (`score novelty or parsimony`). Moves in this case were given a score calculated by subtracting the number of topologically novel nodes from the parsimony change. In this case, moves that achieved higher negative values were considered better. This allowed subparsimonious structures to be included in the MADAG.

We found that allowing moves that improve either parsimony score or topological variation produces a more effective tree search than scoring SPR moves by optimality alone (Fig. 3). We compared the results for 30 replicates using each scoring option at the end of 30 sample-optimize-merge iterations for each case. The results are shown for an alignment containing 245 taxa for dengue fever obtained from the Atlas of Viral Adaptation Evolution project [17].

**Figure 3:**
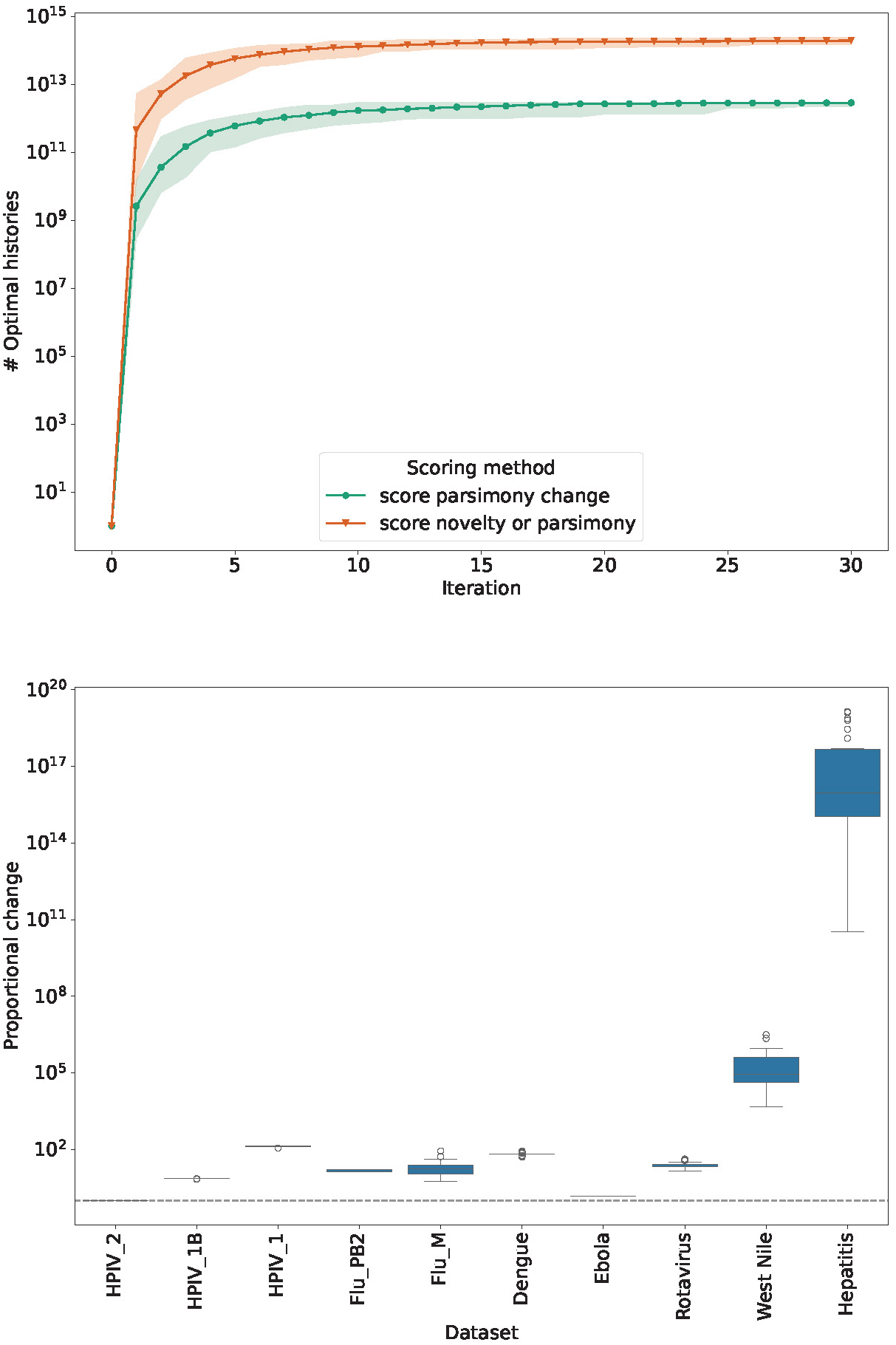
Comparison of topology-based and parsimony-based criteria to evaluate SPR moves in `larch`. The top plot shows accumulation of parsimony-optimal histories in the MADAG over 30 sample-optimize-merge iterations for the `Dengue` dataset (30 replicates; shaded regions show range, solid lines show means). `score parsimony change` scores moves by parsimony improvement; `score novelty or parsimony` scores by the sum of parsimony and topological novelty. The bottom plot shows the distribution of proportional increase of history count from using `score novelty or parsimony` relative to `score parsimony change` (dotted line indicates identity).

The average number of optimal histories found using the scoring method that incorporates topological novelty was greater in general than that found using the parsimony criterion, with a median proportional increase of approximately 20.658.

### Sampling novel starting topologies accelerates search for optimal histories

In addition to the choice of which moves are accepted, another feature that we used to guide tree search is the choice of sampled history on which to calculate SPR moves. We considered five probability distributions for sampling a history as the starting point for each sample-optimize-merge iteration. The first distribution, `sample MP history`, is the natural distribution on the optimal histories in the MADAG. This distribution is defined by assigning equal weight to each edge descending from a given clade. The second distribution, `sample any history`, is the natural distribution on the unrestricted set of all histories in the MADAG. The third distribution, `sample medoid history`, samples from the natural distribution on the set of histories in the MADAG that realize a minimum summed RF distance to the other histories in the MADAG. That is, these are the histories that are close to the topological centroid of the MADAG. The fourth distribution, `sample novel history`, samples from the natural distribution on the set of histories that maximize the summed RF distance over all of the histories in the MADAG. The final distribution, `sample uniformly`, samples uniformly from the set of all histories in the MADAG (we mention this distribution for completeness though don’t use it in the experiments here).

Starting with a single history, we compared the results of running `larch` using each of the sampling methods on the same datasets that we used to evaluate scoring methods. For each sampling method, we ran 30 replicates of 30 sample-optimize-merge iterations.

Sampling topologically novel histories produced more MP histories than sampling optimal histories overall (Figs. 4 and 5). In a subset of cases, sampling a medoid history performed similarly to sampling a novel history. This may be due to the fact that the space of histories in these cases is not shaped like a ball, and so a medoid history is in fact not the center of a dense neighborhood of histories. We further explore the shape of the space of histories based on summed and pairwise RF distances later in the paper.

**Figure 4:**
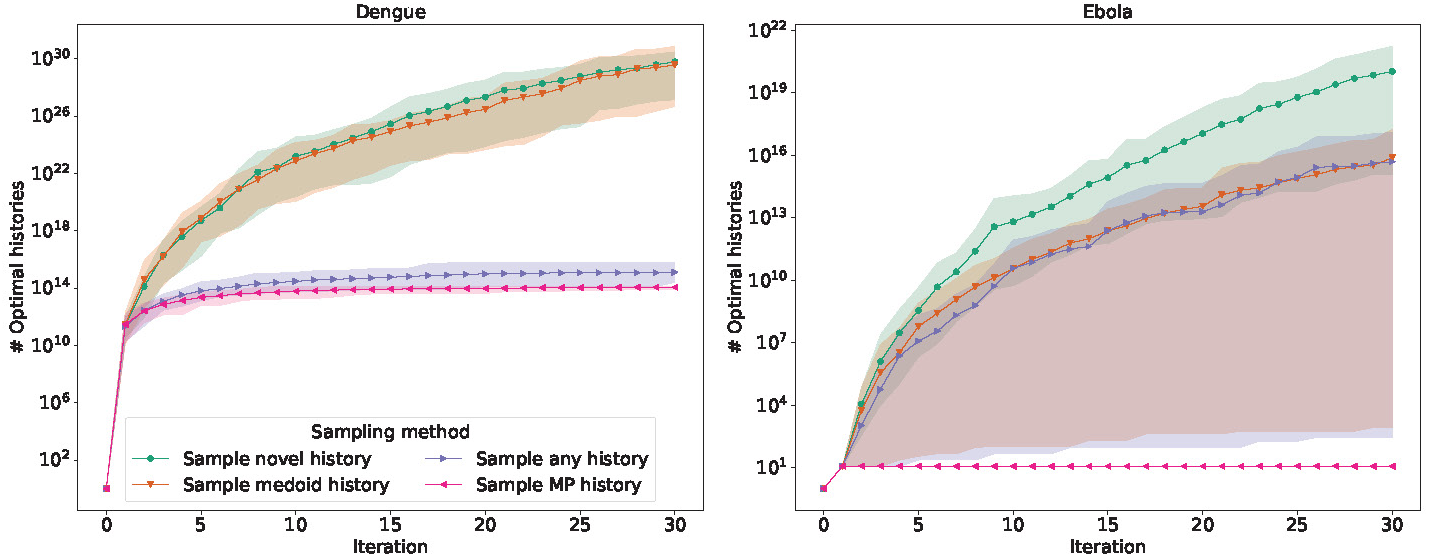
Comparison of sampling distributions for histories from the MADAG across two datasets (Dengue: 245 sequences; Ebola: 297 sequences). Accumulation of parsimony-optimal histories over 30 sample-optimize-merge iterations (30 replicates; shaded regions show range, solid lines show means). `sample MP history` samples optimal histories; `sample any history` samples any history; `sample medoid history` and `sample novel history` sample topologically mean or novel histories, respectively.

**Figure 5:**
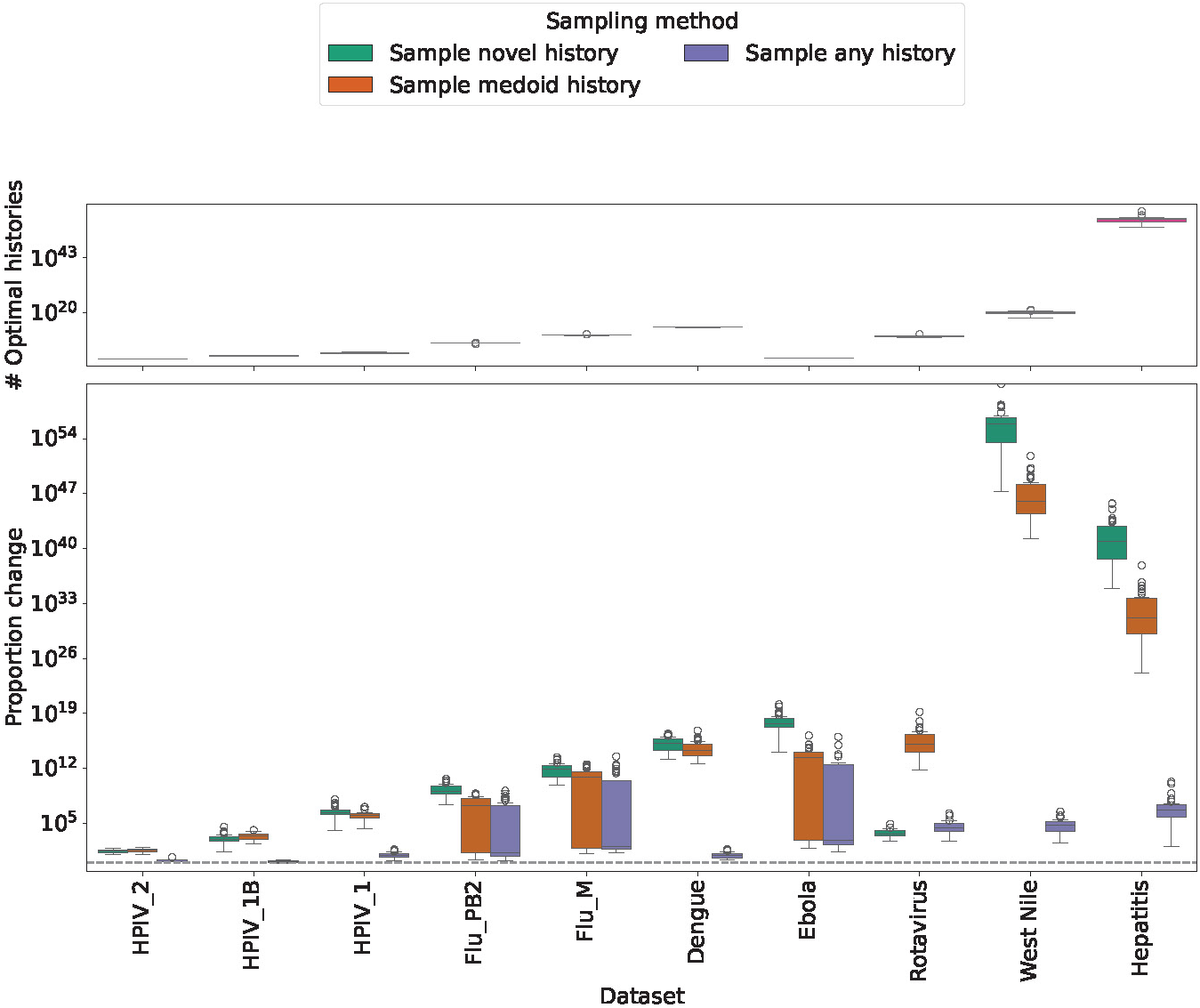
Comparison of sampling distributions for histories from the MADAG across datasets (ordered by MSA length). Top plot shows final parsimony-optimal history counts after 30 iterations using `sample MP history`. Bottom plot shows proportional increase/decrease of alternative sampling methods relative to `sample MP history` (computed by dividing by the mean of the `sample MP history` distribution).

Contrary to our expectation, we found that sampling potentially suboptimal histories as starting points for SPR search produces significantly more optimal histories than using strictly optimal histories. Indeed, we conducted the same experiment across all 10 viral datasets under consideration. Across the majority of datasets, the sampling method `sample novel history` produced the largest number of optimal histories (Fig. 5). This shows that rewarding topological novelty allows the SPR search to escape local optima to explore a wider range of optimal histories.

### Maintaining ambiguities preserves history diversity

We also found that preserving original ambiguities in the data improved tree exploration. The `larch` framework has been extended to admit sequence data that contains ambiguities on leaf nodes. The optimization process in `matOptimize` requires a disambiguated history. To deal with this, the `larch` implementation includes an additional step where the sampled history is disambiguated before being used for the SPR search. The mutations on pendant edges are therefore fully disambiguated, as they are inherited from the fully disambiguated MAT structures that are merged in.

There is only a small variation in outcome between using ambiguities and not using them when compared across a range of starting topologies (Fig. 6). However, preserving the ambiguities in the data is necessary when the initial starting history is suboptimal. We compared the results of `larch` using ambiguous versus disambiguated data for a subset of the datasets which contain ambiguities. For each dataset, we used a near-optimal starting history and used Sankoff to compute optimal disambiguations for the data based on those histories. We ran 30 replicates for each starting history and each option, and compared the number of optimal histories that were found after 30 sample-optimize-merge iterations.

**Figure 6:**
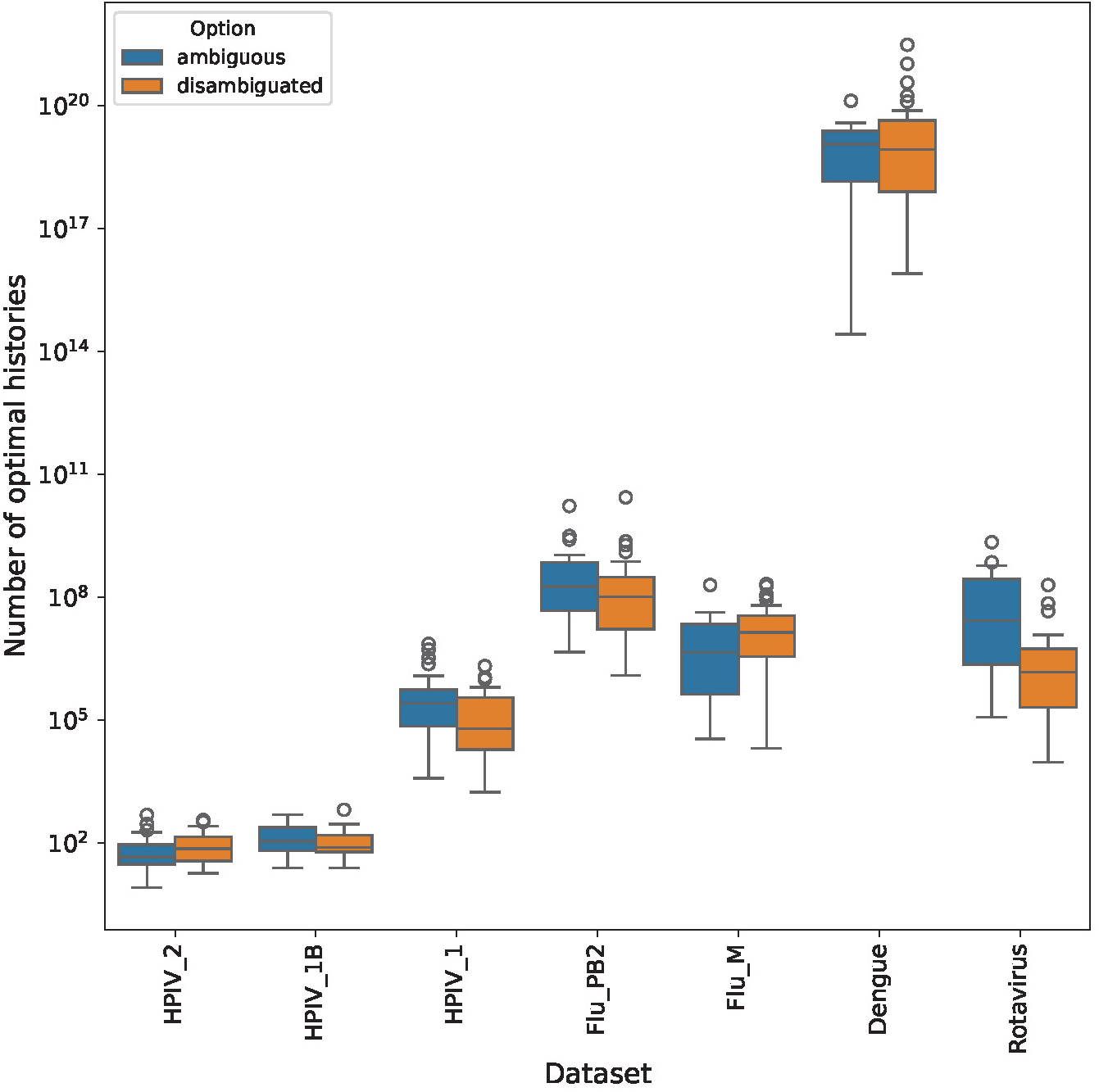
Preserving sequence ambiguities leads to similar numbers of optimal histories compared to disambiguated data. Comparison of optimal history counts for ambiguous and disambiguated data.

The datasets were ordered by size, with the shortest MSA on the left. In the case of `HPIV 2`, only one optimal topology was found by either method. The set of optimal histories differs only in the resolution of Fitch sets.

In order to maintain parsimony optimality for all histories in a MADAG, we require that all internal nodes have fully disambiguated data. This is necessary because nodes can have multiple parents as well as multiple children in the MADAG, and hence maximally parsimonious resolutions of an ambiguous base are not always compatible.

### The MADAG reveals the structure of optimal history space

Using the MADAG, we characterized the optimal history space for each dataset. To summarize our key conclusions, we found:

1. well-defined clusters of topologically distinct but <u>equally parsimonious</u> histories, which we will call “plateaux”
2. extremely large topological differences between these plateaux for some datasets
3. different datasets show different distances, configurations, and densities for the plateaux

We will describe these results after giving a brief description of the analyses that led to these conclusions.

The MADAG is a natural instrument for storing large ensembles of parsimonyoptimal histories. We implemented dynamic algorithms to compute the RF distances for all histories in the MADAG. These algorithms effectively calculate the pairwise and summed RF distances between a given history and the rest by traversing the edges of the MADAG. This allows us to compute the sum of RF distances between each history in the MADAG and the others, as well as the maximum and minimum.

We used these algorithms to calculate lower and upper bounds on the diameter of the space of histories in the MADAG. These bounds give an approximation of the amount of topological diversity in the space of histories represented by the MADAG. The first method provides a lower bound on the true diameter by finding a maximum distance from a specific history. Specifically, it samples a topologically novel history, which is a history that achieves maximal summed RF distances over all histories in the MADAG. This topologically novel history is then used to calculate the maximal RF distance between it and all other histories in the MADAG. The second method, which is an upper bound, computes an approximate radius and doubles it. The approximate radius is calculated by sampling a “medoid” history, which minimizes the summed RF distance over all other histories in the MADAG. The upper bound is found by doubling the maximum distance between the medoid history and any other history. This is indeed a true upper bound, since the triangle inequality gives that the maximum RF distance between any two histories is bounded by twice the distance between any history and the medoid.

These upper and lower bound calculations together with the pairwise RF distances provided summary information about the space of optimal histories, which we combined with a comparison of distributions of RF distances for insight into the grouping of optimal histories as well as the overall shape of optimal tree space.

The RF bounds show an extremely large diameter over the set of optimal histories for several datasets (Fig. 7). A dataset with high RF bounds and a small average pairwise RF distance has many highly similar histories that are close to each other in terms of RF distance, but they are likely clustered into tightly packed groups. On the other hand, a dataset with an average pairwise RF distance that is close to the lower bound on RF diameter likely has disparate optimal histories, with little or no clustering.

**Figure 7:**
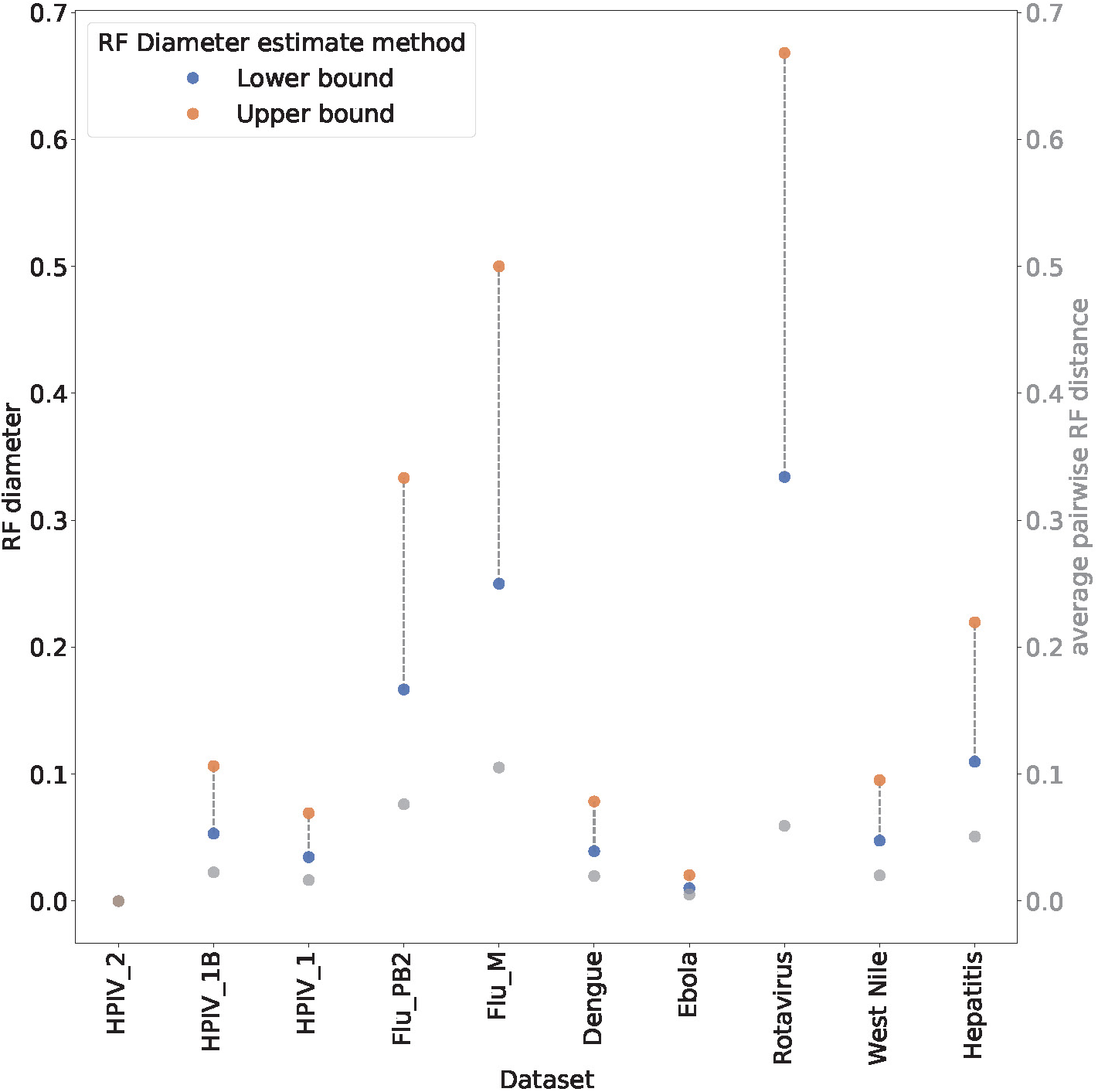
Lower and upper bounds on the approximate diameter of the optimal history space show that there are often pairs of highly divergent MP histories. Average pairwise RF distance over all optimal histories is also shown with gray points. All RF distances are normalized by the maximum possible RF distance.

We sought to explore the variety and diversity of topologies in the space of optimal histories. The average pairwise RF distance does not provide insight into the underlying distribution of RF distances among optimal histories, and so we performed the following additional investigation. We computed for each history in the MADAG the sum of pairwise RF distances to all other histories in the MADAG. This gives the range of summed RF values that are achieved and shows how they are grouped. We also computed the pairwise RF distances from a medoid history to all of the remaining histories (we call this the “medoid distribution”) and the pairwise RF distances from a topologically novel history to the remaining histories (we call this the “outlier distribution”). The distribution of pairwise RF distances to a specific topology gives a sense of how the histories are grouped with respect to that specific history. We took two topologically meaningful points in the space of histories to gain a broader understanding of the space using pairwise distances.

Using a combination of the RF diameter bounds and the tree RF distributions, we found a wide variety of tree spaces, including spaces with multiple welldefined clusters of similar topologies (Fig. 8, with additional results in Fig. S2). These were notably different both in terms of the tightness of the clustering and in terms of the number of plateaux or clusters that were represented in each space. We now describe those in detail.

**Figure 8:**
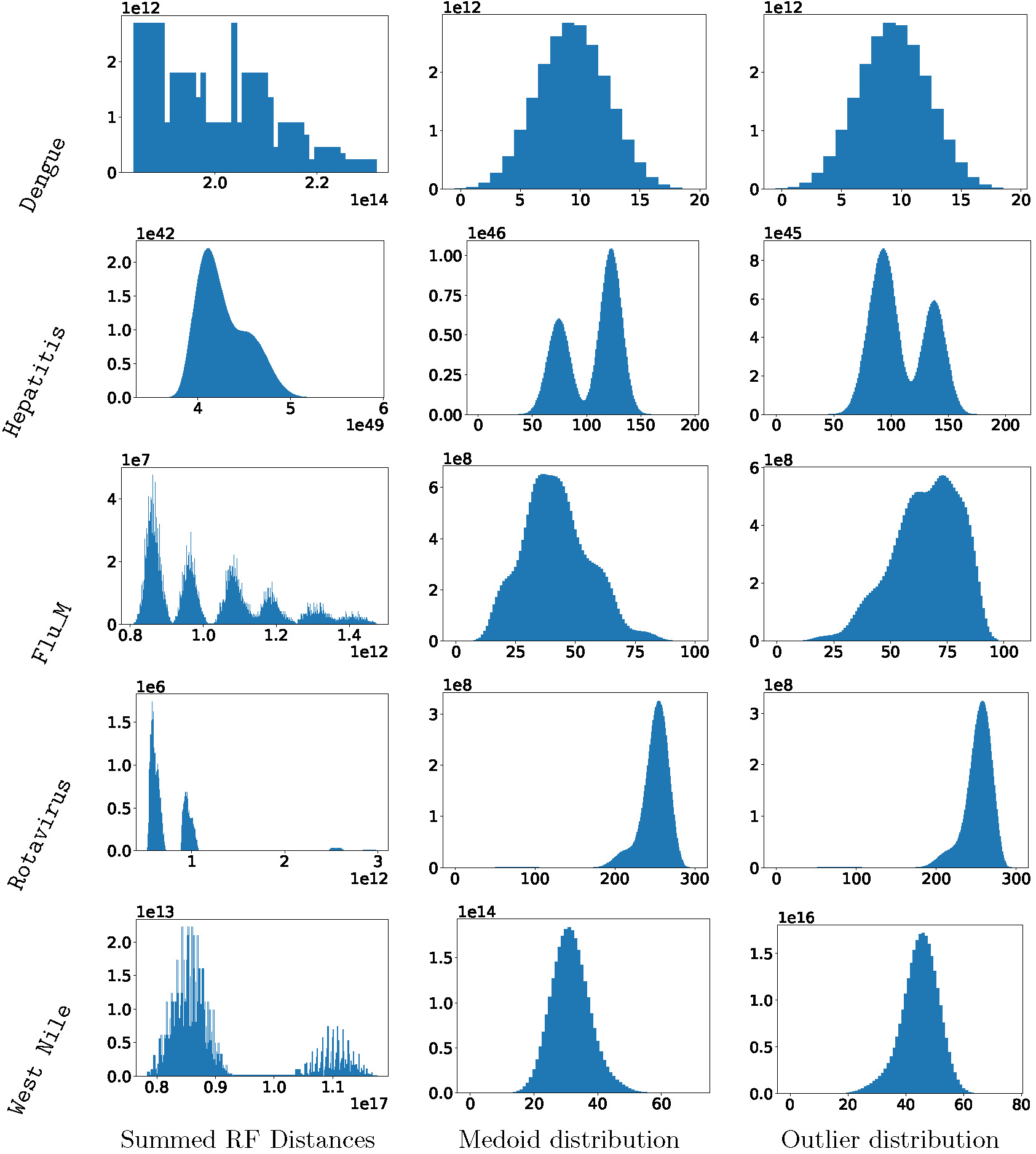
Diverse structuring of optimal histories across datasets (each row = one dataset). Left: summed RF distances for all histories. Center: pairwise RF distances to medoid history. Right: pairwise RF distances to novel history. Y-axis shows counts.

#### Single plateau space

The first kind of space is a collection of optimal histories whose topologies are not clustered into multiple distinct groups. We see for the `HPIV 2` dataset that the upper and lower bounds are both 0, thus, this set of optimal histories contains a single unique topology. More interestingly, the remaining `HPIV` datasets as well as `Dengue` and `Ebola` have small values for all three distances. This indicates that the optimal histories are topologically similar. In these cases, we can assume that the space of optimal histories represented by each MADAG contains a single cluster or “plateau” of optimal histories. The `Flu PB2` dataset shows a single peak, however, the asymmetric tails indicate that there might be a second smaller peak contributing.

#### Multiple plateaux

The `Rotavirus`, `Hepatitis`, and `Flu M` datasets have multiple plateaux, exhibiting small average pairwise RF distance when compared with the upper and lower bounds (Fig. 7). Furthermore, in all of these cases there is a significant difference between the bounds. That is, the maximum distance between a topological outlier and any other history is significantly smaller than twice the distance between that outlier and the medoid. This implies that the space of optimal histories has two or more plateaux, but there is one mode that is significantly farther from the medoid than any other. Thus the space is not star shaped, since there are not multiple clusters of distinct and equally distant topologies.

The distribution of distances allows us to understand the shapes of these sets of optimal histories (Fig. 8). The plot containing summed RF distances for all histories on the `Rotavirus` dataset shows two distinct peaks, each with an asymmetric distribution. Recall that the medoid is the tree that minimizes the summed RF distance over all of the histories. It can be seen from the plot for summed RF distance that in many datasets there is a well-defined peak in the neighborhood of the smallest summed RF distance value, indicating a cluster of optimal histories close to the medoid history. The farthest peak indicates that there is at least one cluster containing the most topologically novel history, and it would seem from the dual peak structures that there is another cluster of histories that is closer to the medoid but distinct from it.

In the case of `Flu M`, we see multiple peaks in the distribution of summed RF distances, as well as in the medoid distribution and the outlier distribution. This implies a diverse set of topologies with 5 or 6 distinct plateaux. Both the `Hepatitis` and `West Nile` datasets show two distinct peaks, indicating two plateaux in the space of optimal histories.

We also investigated the density of each cluster. Both `Flu M` and `Hepatitis` datasets show well-defined peaks in the plot. The peaks in the medoid and outlier distributions for `Hepatitis` are nearly disjoint, indicating that the two clusters are well-defined and distinct. However, the RF diameter for this dataset is small, showing that these two plateaux contain histories tightly clustered around two different backbone topologies that share a significant proportion of their splits. In contrast, the `Flu M` dataset has an extremely large RF diameter. The space of optimal histories for this dataset has well-defined peaks indicating clusters centered around widely differing topologies.

### Different starting histories yield similar results

A common difficulty with tree search methods is how to explore regions beyond a neighborhood of the initial starting point; here, we sought to understand to what extent `larch` was exploring the entire space versus getting stuck in local optima. In other contexts [7, 8] we have found it informative to start search in multiple places, and investigate the degree to which these multiple starting points converge to the same result. For the current MADAG context, we can take multiple starting histories, put them in a DAG, and compare that to exploration using a single starting history. If `larch` has limited ability to explore, the multiple starting history approach should result in a larger set of histories than the single starting history approach.

Instead, we recovered a similar number of histories using a single starting history and using a MADAG built from multiple different histories (Fig. 9). For this analysis, we initialized four different starting histories using UShER [14] and compared the results after 30 iterations with the results from using a starting point of a MADAG built from the four initial topologies. The sequences were randomly shuffled before creating each UShER history.

**Figure 9:**
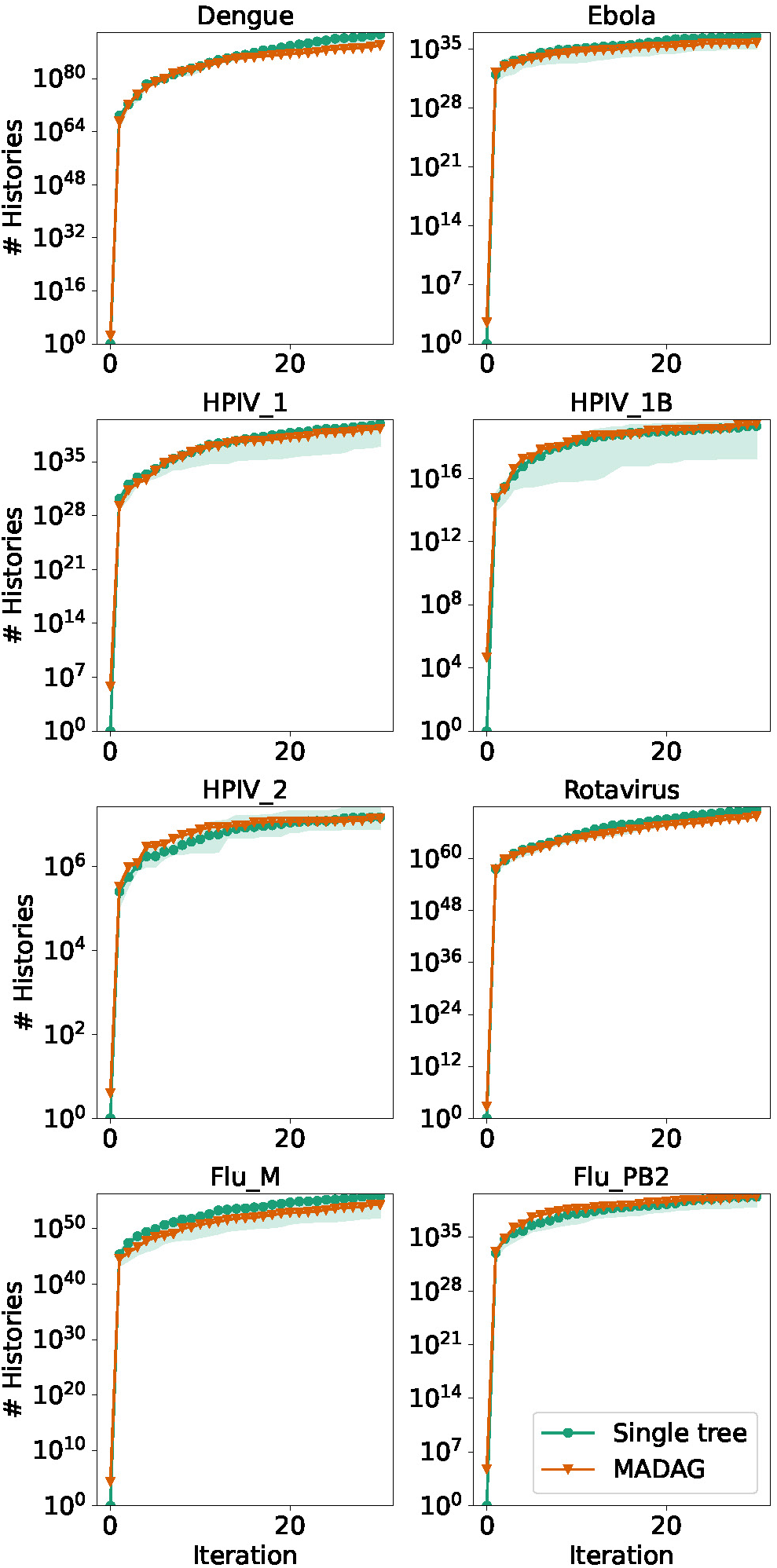
Number of histories found at each iteration when running `larch` with a starting MADAG built from 4 UShER starting histories compared to the number of histories found with a single starting history. The shaded region denotes the range of values found using four different starting histories. The solid line corresponds to the average value of the history counts for each of the four parallel runs at each iteration.

### larch produces highly optimal trees

The `larch` software is guaranteed to produce a history with the same or better parsimony score than `matOptimize`. Using `matOptimize` in the search phase of `larch` ensures that the best results found by `matOptimize` are included in the output. An example of this parsimony improvement is shown in Fig. S1 for the `Dengue` dataset. For all datasets considered in this study, the results showed either identical parsimony scores to those found by `matOptimize` or slight improvements when using `larch`.

### larch is memory efficient and computationally efficient

The MADAG structure allows `larch` to represent a vast number of histories in compact form, enabling execution on a standard laptop computer. It is also highly optimized to explore a large number of histories in a short time. The results for 10 iterations of `larch` on each dataset using a single starting history and default parameters are given in Table 1 on a device running Ubuntu 22.04 with Intel Core i7 Processor(4x 5GHz) and 32 GB LPDDR RAM.

**Table 1:** Results of running 10 iterations of larch with default parameters for datasets of differing sizes. Values for runtime duration are given in seconds, and values for peak runtime memory usage are given in GB.

| Dataset | # Sequences | Runtime (s) | Peak usage (GB) | # Histories |
| --- | --- | --- | --- | --- |
| Dengue | 245 | 115.87 | 2.453 | $5.0391 \times 10^{57}$ |
| Ebola | 297 | 397.92 | 4.254 | $3.5200 \times 10^2$ |
| Hepatitis | 1,032 | 7,944.65 | 28.01 | $1.6726 \times 10^{285}$ |
| Flu_M | 215 | 247.9 | 3.78 | $3.4293 \times 10^{22}$ |
| Flu_PB2 | 135 | 36.59 | 1.817 | $1.2383 \times 10^{23}$ |
| HPIV_1 | 104 | 25.26 | 2.364 | $2.0406 \times 10^{27}$ |
| HPIV_1B | 50 | 9.44 | 1.8 | $2.7418 \times 10^{13}$ |
| HPIV_2 | 19 | 5.06 | 2.481 | $7.1344 \times 10^4$ |
| Rotavirus | 452 | 1,060.11 | 8.252 | $5.7142 \times 10^{21}$ |
| West Nile | 801 | 5,619.36 | 26.982 | $3.6506 \times 10^{130}$ |

## Discussion

In this work, we introduce the MADAG, a data structure that combines the compact mutation-level edge annotations of the MAT with the DAG structure’s ability to represent large families of trees and histories. We implement the MADAG in `larch`, a highly optimized C++ program that includes novel algorithms for parallelized fragment merging, dynamic programming on DAGs for computing RF distances and sampling histories, and an overlay structure that enables lightweight storage and application of SPR moves to the MADAG. These technical capabilities enable new approaches to phylogenetic tree search.

Unlike existing methods that optimize single trees, `larch`’s ensemble approach enables valley-crossing exploration that <u>systematically</u> maps optimal tree space structure. This is different from previous methods that work to cross valleys using perturbations of the objective. For example, the ratchet [18, 19] repeatedly bootstraps data to break out of local minima. For Bayesian analyses, the MC^3^ algorithm uses a combination of parallel Markov chains with different temperatures to allow some chains to wander more freely [20, 21]. In contrast, `larch` uses the entire ensemble of optimal histories to direct the search under the original objective. This is effective: we show that `larch` does better than `matOptimize` for finding the best histories, and `matOptimize` in turn was shown to outperform TNT [22], which uses the ratchet, sectorial search, tree drifting, and tree fusing.

`larch` also integrates ambiguities in the data, which is more technically challenging in the DAG case than for single trees. Ambiguities are necessary for ensuring that all possible optimal reconstructions are considered. Indeed, we showed that in some cases, using ambiguities allowed `larch` to identify histories with a strictly better parsimony score than was found using disambiguated data.

We have explored the vast space of optimal histories for a range of datasets, and found that it has complex structure. Specifically, the optimal tree space often contains multiple clusters, indicating topological diversity that is fully represented in the MADAG. This is enabled by dynamic programming on the DAG that allows us to extract summary information about the space of optimal histories, including the formation and grouping of clusters in optimal history space.

We believe that the `larch` approach is successful while an analogous approach for likelihood-based analysis [12] was not because every history in the MADAG after trimming is optimal. In contrast, the sDAG used by [12] may include many suboptimal trees. Moving forward, it would be interesting to use `larch` and the MADAG as a means of exploring tree space under a likelihood-based criterion.

Stepping back, we view the MADAG as part of the evolution of compact phylogenetic data structures with roots in [15] that exploit the low diversity of densely sampled molecular sequences. The MAT [14] pioneered mutation-level edge annotations for single histories, storing only a reference sequence at the root rather than full sequences at every node. More recently, Delphy [23] introduced Explicit Mutation-Annotated Trees (EMATs) for Bayesian phylogenetics, which similarly represent sequences implicitly through accumulated mutations from an explicit root sequence. However, unlike these previous efforts, the MADAG is able to capture large families of optimal histories with full optimality information and mutation-level edge annotations in one data structure, enabling new methods for systematic exploration and a new way of describing the collection of optimal histories.

## Materials and methods

### Data

We used a set of 11 viral datasets that are listed with full names and sizes in Table 2. We obtained 8 datasets from the Atlas of Viral Adaptation project hosted on https://github.com/blab/adaptive-evolution. The 20B dataset is taken from the clade of the same name from the UShER history reconstruction for SARS-CoV-2 [24]. The final 2 datasets were for Ebola virus and West Nile virus, obtained from separate epidemiological studies [25, 26].

**Table 2:**
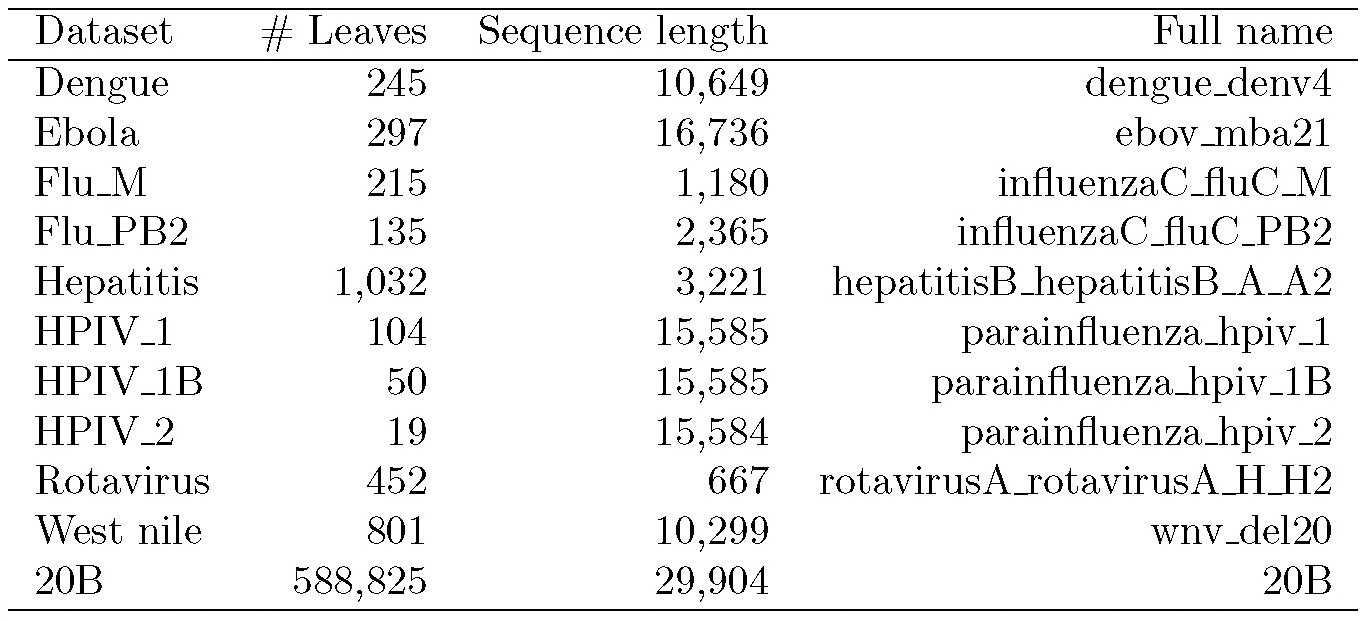
Viral datasets used in this study.

### Lightweight storage implementation

#### The MADAG data structure

The MADAG storage structure is a form of directed acyclic graph in which a full sequence is stored at the root node. Each edge is annotated with the set of mutations corresponding to the site and value where the parent and child nodes’ sequence data differ. For example, if the two sequences contain different bases at site ten, with the parent sequence having base “A” and the child sequence base “C” at that site, then the mutation set consists of the single mutation “A10C.” In the densely sampled regime, where sequence lengths are thousands of characters long and the parent and child sequences in such histories typically differ by one to two mutations, this provides a massive reduction in memory use. Additionally, each node is annotated with a compact genome, which is the set of mutations for the given node relative to the reference sequence.

#### Overlay for SPR moves

A major computational burden of the sample-optimize-merge structure is storing and applying SPR moves to the MADAG. In order to fully recreate an SPR move, we require a source node, a target node, the set of nodes along the path between the source and target nodes, and any node ancestral to the SPR path which has a changed Fitch set after applying the move and running the Fitch algorithm. In order to avoid creating an entirely new tree-shaped structure for each SPR move, we implemented an overlay class that interfaces with the sampled history being used for SPR search, but which has redirected access for the node and edge connectivity structure as well as for edge mutations. This lightweight class allows us to store the changes relative to the sampled history. Each SPR move is stored as an overlay object which covers the connected subset of the sampled history containing the nodes that would be altered by the given SPR move together with any parent or child edges of those altered nodes. We call these overlaid substructures “fragments.” We implemented a highly parallelized algorithm to merge collections of fragments into the MADAG concurrently. During the SPR search phase, each move that is approved is added to a bucket, and when the bucket reaches capacity, these moves are merged directly into the MADAG. This ensures that memory usage is controlled while at the same time optimizing parallelized fragment merging.

### Dynamic programming on the MADAG

#### Calculating RF distances

We compute Robinson-Foulds distances between a given history and the remaining histories using a dynamic program. The algorithm computes the number of histories in which each node takes part in an initial traversal of the MADAG. This cached data is then used in combination with the clades to calculate the RF distance between any reference history and the remaining histories in the MADAG. The minimum, maximum, and summed RF distances can be calculated by aggregating the data using the corresponding mathematical operations at each clade in a single traversal of the edges of the MADAG.

#### Sampling histories

Histories are sampled from the MADAG in a preorder traversal beginning at the root node. For the child node of each edge that is chosen we specify a single edge descending from each of the node’s clades. Assigning numeric weights to the edges in the MADAG induces a corresponding probability distribution on the set of histories that it represents. We implemented a dynamic program that annotates the edges of the MADAG with weights to induce a range of discrete probability distributions including a uniform distribution, and parsimony and topology-weighted distributions.

The main feature contributing to runtime cost for `larch` is the fact that each possible SPR move is evaluated. In order to optimize more locally, we implemented an option to sample subtree structures as an alternative to sampling histories constructed on the entire leaf set. This option can be used to guide the optimization search to focus on substructures if the larger optimal backbone topologies have been identified.

## Acknowledgements

We would like to thank Hugh Haddox and Angie Hinrichs for testing larch.

This work was supported by NIH grants R01-AI146028 and R01-AI162611. Scientific Computing Infrastructure at Fred Hutch is funded by ORIP grant S10OD028685. Frederick Matsen is an investigator of the Howard Hughes Medical Institute.

## Supplementary Materials

Supplementary material, including data files, can be found in the Zenodo data repository DOI 10.5281/zenodo.17467197.

### Additional figures

**Figure S1:**
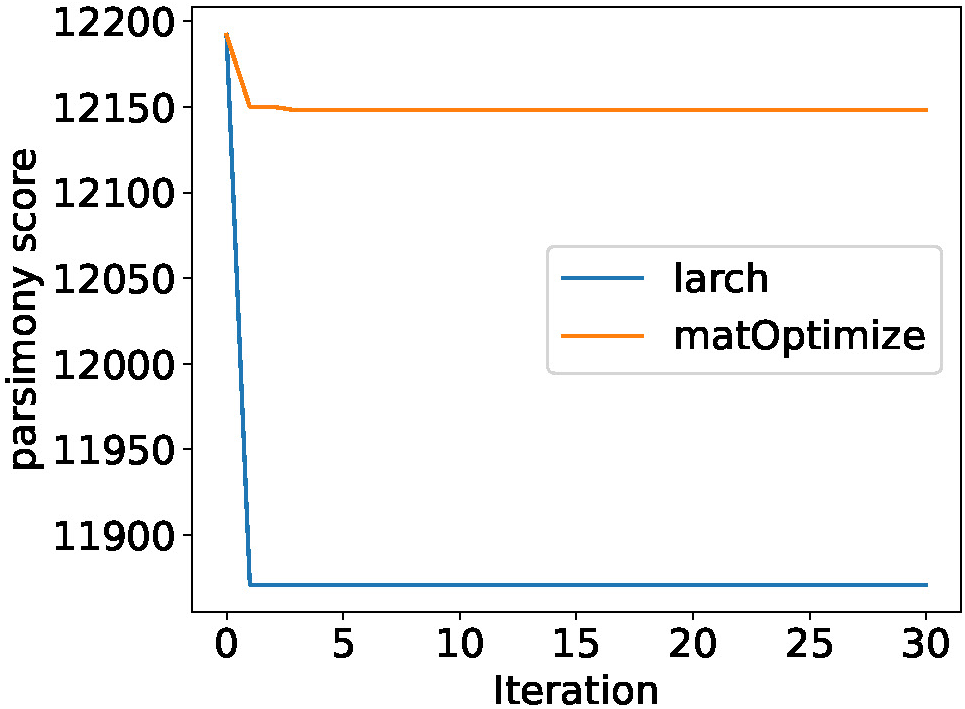
Comparison of the best parsimony score found by `larch` and `matOptimize` at each iteration for the Dengue dataset. The same starting history was used for each case.

**Figure S2:**
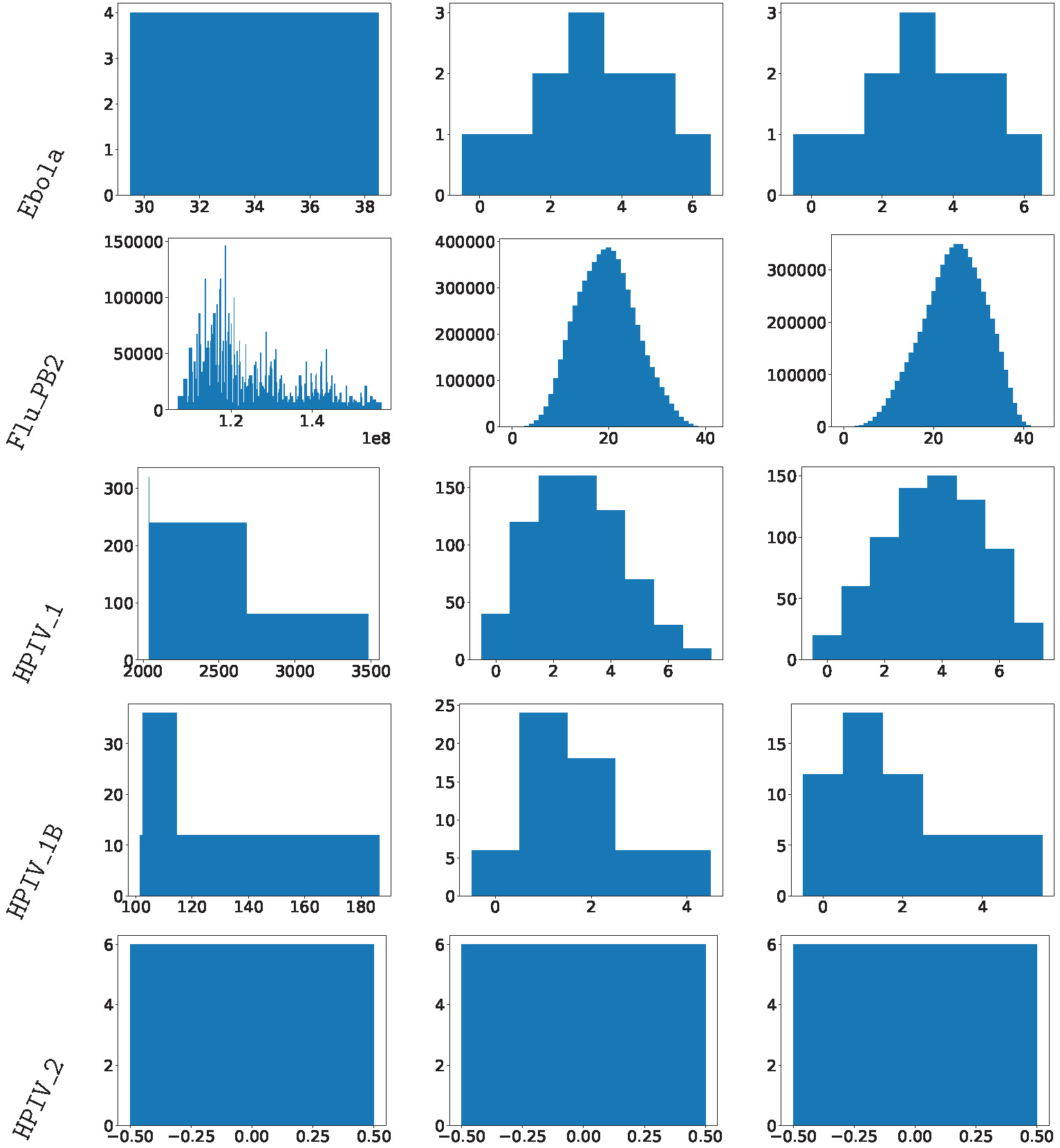
RF distance distributions reveal diverse clustering patterns in optimal history space across additional datasets (each row = one dataset). Left: summed RF distances for all histories. Center: pairwise RF distances to medoid history. Right: pairwise RF distances to novel history.

### larch flags

There are a variety of optional arguments that can be used to optimize tree search using `larch`. For example, the option `move-coeff-nodes` can be used to bias the search toward SPR moves that introduce more novel features. This option specifies the factor that multiplies the number of new nodes each SPR move introduces. A factor of 0 causes `larch` to only accept SPR moves that preserve or improve parsimony. A factor of 1 or greater causes `larch` to incorporate the scaled number of new nodes into the score.

**Table S1:** Optional parameters that can be used to customize the search.

| Option name | Description |
| --- | --- |
| **--autodetect-stoptime** | Terminate once the best parsimony score found at each iteration reaches a plateau. |
| **--keep-fragment-uncollapsed** | Keep edges that have no mutations associated to them, rather than collapsing them. |
| **--min-subtree-clade-size** | The maximum number of leaves in a subtree sampled for optimization. |
| **--max-subtree-clade-size** | The minimum number of leaves in a subtree sampled for optimization. |
| **--max-time** | Exit after fixed runtime(in minutes). |
| **--move-coeff-nodes** | Specify a scaling factor for how the number of new nodes contributes to SPR move score. |
| **--move-coeff-pscore** | Specify a scaling factor for how the change in parsimony contributes to SPR move score. |
| **--sample-method** | Select distribution to use for choosing a sampled history. |
| **--switch-subtrees** | Switch to optimizing subtrees instead of full trees after the specified number of iterations. |
| **--trim** | Trim the final MADAG to contain only maximally parsimonious histories. |

